# Brainacle: Interactive delineation and quantification of anatomical structures with virtual reality

**DOI:** 10.1101/2025.06.17.659041

**Authors:** Dimitar Garkov, Etienne Lein, Niklas Kielkopf, Christian Dullin, Karsten Klein, Bjorn Sommer, Alex Jordan, Falk Schreiber

**Affiliations:** Department of Computer and Information Science, University of Konstanz, Konstanz, Germany; Behavioural Evolution Research Group, Max Planck Institute of Animal Behavior, Konstanz, Germany; Department of Biology, University of Konstanz, Konstanz, Germany; Translational Molecular Imaging, Max Planck Institute for Multidisciplinary Sciences, Göttingen, Germany; Elettra Sincrotrone Trieste ScpA, Trieste, Italy; Institute for Clinical and Interventional Radiology, University Medical Center Göttingen, Göttingen, Germany; School of Design, Royal College of Art, London, United Kingdom; Faculty of Information Technologies, Monash University, Melbourne, Australia

## Abstract

A virtual environment, equipped with suitable data representations, interaction techniques, and method interfaces, can facilitate effective human-in-the-loop analytics of 3D data. Motivated by this observation, we propose *Brainacle*, a native editor for the interactive delineation and quantification of anatomical structures in virtual reality. We validate this in a case study, where we process the gross brain regions of 20 individual animals from six different species of African cichlid fish from the tribe *Lamprologinii*, endemic to Lake Tanganyika in Africa. The software presented in this article can be used to quickly visualise and obtain an overview of imaged animal tissues, visually communicate features of interest, interactively add delineations, and perform different measurements in the same 3D environment. *Brainacle* is made available free of charge at https://brainacle.org and on the research data repository DaRUS.

## Introduction

Consensus is still lacking whether virtual 3D environments (VE) are indeed more expressive than their 2D counterparts. The decision whether or not to use VE, therefore, should be motivated by the specific use case and the distinct benefits the application of those environments could provide [1]. One well-known benefit of VE is how immersion can predispose mental involvement [2], so that attention and time may be perceived differently when in the state of flow [3]. Another benefit VE can offer, first described some 30 years ago, refers to how a user’s performance at certain tasks may be enhanced using various different perceptual overlays [4, 5]. In the context of multiple users, well-designed VE also have the potential to support social learning and synchronous collaboration, thus indirectly helping users gain expertise in the domain.

In parallel, recent advances in biological imaging have enabled high-resolution, whole-tissue imaging. With imaging data that is inherently three-dimensional, one can expect VE would become integrated in more imaging pipelines. These pipelines, however, often involve some form of segmentation, which remains a laborious process in spite of the recent proliferation of AI-based tools. This is because validation requires a human in the loop with expert knowledge, a need that is even more pronounced when the imaged species is less well studied and non-human [6]. And while there has been substantial progress recently, the performance of segmentation models, such as MedSAM 2 [7], still depends on many different parameters and it may not be readily applicable to new, unseen data.

In the current work, we applied the following biological pipeline: imaging → reconstruction → delineation → quantification → analysis to a case study of African cichlid fish from the tribe *Lamprologinii* to study the research question of how to quantify the brain structure of closely related species of divergent social behaviour. Given the high number of imaged animals and tissues, which lacked previous detailed 3D brain atlases, we considered exploring alternative methods, such as virtual reality (VR). Considering the interactivity VR provides, we could specifically target the latter part of the pipeline: delineation and quantification. Our approach to delineation thus does not rely on a specific image processing protocol or involve the use of black-box segmentation models, which may not generalise to other species or data. Instead, we apply VR to derive higher-level structural information from simple user input. To achieve this, we developed *Brainacle*, a native VR editor designed in close collaboration with domain experts. It enables the interactive delineation and quantification of anatomical data inside VR. We make the software freely available at https://brainacle.org and on the data repository DaRUS [8].

While various non-commercial tools and frameworks exist for the analysis of non-human brain tissue images on the desktop [9–12], those targeting VE are considerably less established (one notable exception, although a commercial software tool, being “syGlass” [13]). Among the purely academic, free-to-use VR tools, MiCellAnnGELo [14] has been proposed as a means to annotate segmented time-series microscopy data via surface meshes. vLUME [15] is another VR tool that supports the selection, annotation, and related quantification of point-cloud, single-molecule localisation microscopy data, whose version v1.1 is freely available to academics. Developed in conjunction with 3D Slicer [11], SlicerVR [16] couples interaction, navigation, and collaboration to extend 3D Slicer functionalities to VR. Researchers can also use Diva [17] to view, manipulate, and perform counting and distance measurements on volume renderings in VR, while Usher et al. [18] and TeraVR [19] (part of Vaa3D [12]) enable the tracing of neurons in VR. In contrast to previous design studies, *Brainacle* was used to delineate the gross brain regions of 20 animals from 6 species imaged using synchrotron micro-CT (see Fig. 1 and Tab. 1). Given the variation in social organisation within and across these species, but otherwise remarkable similarities regarding ecology and life history, the study of such datasets, enriched with annotations from the system, may provide new insights into social evolution and its implications on brain evolution [20].

**Fig 1.**
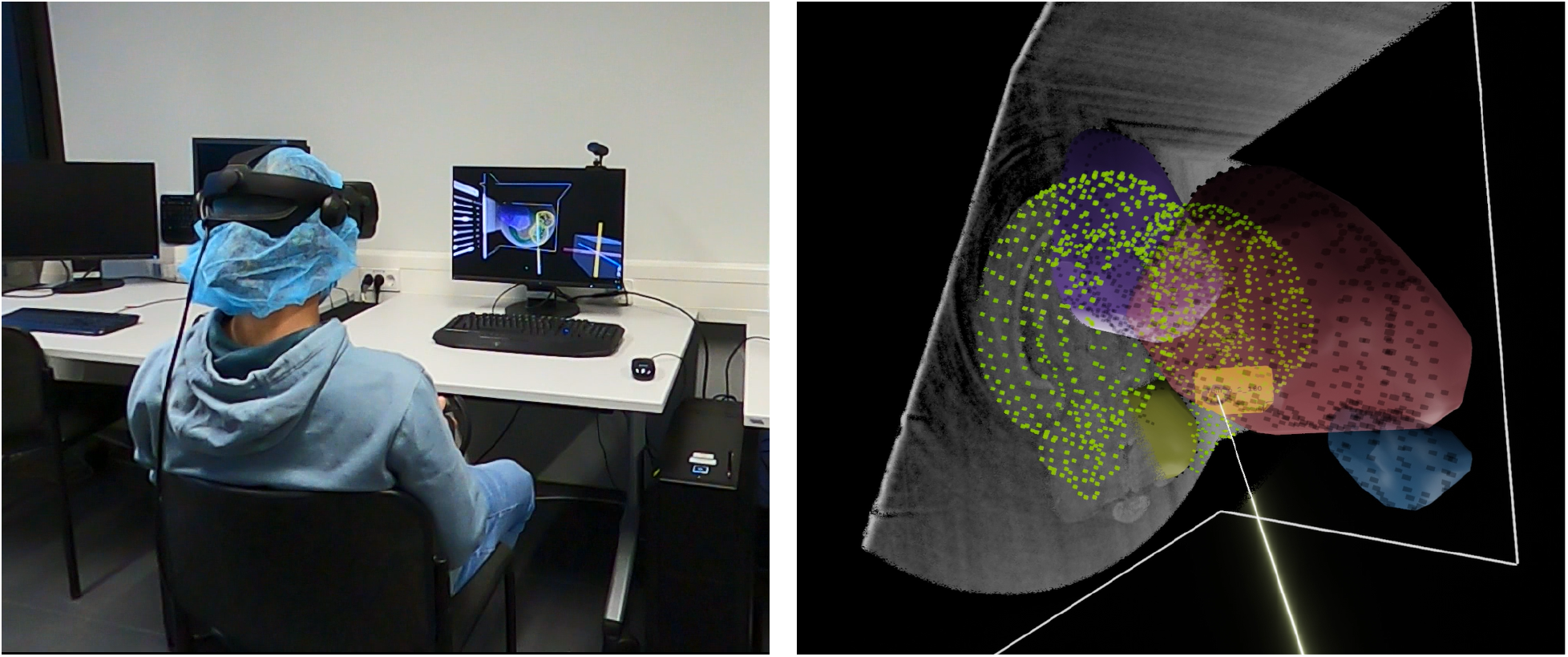
A user obtains an overview of the delineations created with *Brainacle*. A user is shown interacting with the data in VR to gain an overview of the delineations in the brain (left). On the right, a closer view of the delineations can be seen.

**Table 1.** Aggregate measurements per species and gross brain region.

|  | Axis length (mm) |  |  | Volume (mm<sup>3</sup>) |  |
| --- | --- | --- | --- | --- | --- |
|  | Major | Minor | Intermediate | Convex hull | Bounding ellipsoid |
| *Neolamprologus multifasciatus* (N = 5) |  |  |  |  |  |
| **Olfactory bulbs** | 0.708 | 0.364 | 0.518 | 0.039 | 0.071 |
| **Telencephalon** | 1.890 | 1.095 | 1.430 | 1.047 | 1.570 |
| **Optic tectum** | 2.642 | 1.437 | 1.833 | 2.428 | 3.670 |
| **Hypothalamus** | 1.928 | 0.968 | 1.246 | 0.721 | 1.226 |
| **Cerebellum** | 1.743 | 1.329 | 1.455 | 1.114 | 1.803 |
| *Neolamprologus brevis* (N = 2) |  |  |  |  |  |
| **Olfactory bulbs** | 0.931 | 0.439 | 0.633 | 0.073 | 0.136 |
| **Telencephalon** | 2.237 | 1.242 | 1.704 | 1.718 | 2.476 |
| **Optic tectum** | 2.880 | 1.800 | 2.201 | 3.972 | 5.994 |
| **Hypothalamus** | 2.022 | 1.045 | 1.198 | 0.804 | 1.335 |
| **Cerebellum** | 1.762 | 1.467 | 1.520 | 1.260 | 2.064 |
| *Lamprologus ocellatus* (N = 6) |  |  |  |  |  |
| **Olfactory bulbs (5)** | 0.782 | 0.414 | 0.581 | 0.050 | 0.103 |
| **Telencephalon** | 2.101 | 1.299 | 1.738 | 1.680 | 2.572 |
| **Optic tectum** | 3.027 | 1.794 | 2.130 | 4.175 | 6.192 |
| **Hypothalamus** | 2.280 | 1.156 | 1.297 | 1.103 | 1.832 |
| **Cerebellum** | 1.914 | 1.436 | 1.629 | 1.434 | 2.404 |
| *Lamprologus callipterus* (N = 2) |  |  |  |  |  |
| **Olfactory bulbs (1)** | 1.156 | 0.529 | 0.719 | 0.117 | 0.230 |
| **Telencephalon** | 2.210 | 1.505 | 1.741 | 1.905 | 3.067 |
| **Optic tectum** | 3.098 | 1.756 | 2.242 | 4.416 | 6.408 |
| **Hypothalamus** | 2.278 | 1.049 | 1.332 | 1.013 | 1.678 |
| **Cerebellum** | 1.864 | 1.453 | 1.628 | 1.440 | 2.332 |
| *Telmatochromis temporalis* (N = 3) |  |  |  |  |  |
| **Olfactory bulbs** | 0.894 | 0.447 | 0.653 | 0.071 | 0.143 |
| **Telencephalon** | 2.187 | 1.340 | 1.720 | 1.827 | 2.642 |
| **Optic tectum** | 2.927 | 1.696 | 2.284 | 3.779 | 5.938 |
| **Hypothalamus (2)** | 2.243 | 1.106 | 1.431 | 1.189 | 1.861 |
| **Cerebellum** | 2.092 | 1.636 | 1.700 | 1.906 | 3.054 |
| *Lamprologus ornatipinnis* (N = 2) |  |  |  |  |  |
| **Olfactory bulbs** | 0.778 | 0.445 | 0.623 | 0.066 | 0.113 |
| **Telencephalon** | 2.356 | 1.415 | 1.980 | 2.454 | 3.461 |
| **Optic tectum** | 3.382 | 1.972 | 2.356 | 5.452 | 8.227 |
| **Hypothalamus** | 2.245 | 1.272 | 1.369 | 1.130 | 2.049 |
| **Cerebellum** | 2.084 | 1.630 | 1.771 | 1.859 | 3.155 |

### Intended workflow

The user supplies image data, including a voxel size, for visualisation in VR. Delineation is performed by placing geometric primitives over the visualisation in 3D space. Prior delineations can be imported from frameworks such as Fiji and ImageJ [9, 10], as long as all coordinates originate from the same image space. Once delineation is complete, structural features are measured and shown to the user. Computable measurements include counts, distances and volumes, as well as others derived from those. Results are finally stored or exported, allowing visual analysis to be resumed, shared, or versioned.

## Design and implementation

### Data visualisation

In *Brainacle*, imaged data may be visualised either as a direct volume rendering or as an isosurface mesh depending on the data’s preprocessing.

#### Volume rendering

Volume data is rendered 1) using three orthogonal planes or 2) as a sliceable volume rendering. Slicing is used to show exactly the voxels of interest that intersect with the planes in the former case and with a rigid 3D shape in the latter case. This allows each plane to serve as a canvas for delineation that can be shifted while orthogonality is maintained (the 3D shape is not constrained spatially). An accompanying menu allows users to execute several image and volume operations, such as thresholding. Volume data can be imported using the NifTI data format (.*nii*), which is interoperable with software such as ImageJ [9] and Fiji [10].

#### Isosurface mesh

Isosurface meshes are created using image processing software [9, 10] by applying a series of image operations to obtain binary images from which the isosurface mesh is created. The resulting mesh can then be imported in *Brainacle* as a triangle mesh (a Wavefront file). Visibility from both the interior and the exterior of the mesh, implemented using a second shader pass, ensures contouring from both the inside and the outside of the mesh. The interior of the mesh can, for example, be used to quickly measure anatomical ratios and other shape metrics interactively.

### Delineation of structural features

Delineation is a two-step process: 1) generate an input of points (or higher-dimensional simplices), and 2) reconstruct the underlying surfaces from step 1). As such, it is an instance of the problem of surface reconstruction from unorganised points [21]. In fact, all three descriptors—surfaces, contours, and points—form a hierarchy [22, p. 3], which is relevant for analysis later. Hoppe et al. [21] summarised the original problem as: “reconstruct the 3D structures from the stacks of 2D contours”, with algorithms optimised to operate polygon-wise on the sequence of images. To implement interactive delineation in *Brainacle*, we relied on a tool metaphor in order to design two separate tools for delineation. While each of the tools has been designed for stand-alone use, *tool mixing* is also possible, should different substructures part of the same overall structure require different delineation tools.

#### Point tool

The first tool is proposed to enable placing points in a rapid succession, that has been previously seen in mesh painting [14, 16]. We call this the point tool. To use the point tool, the user presses a button to initiate placement. After a short time interval a new point is placed where the tool’s ray is currently casting (from the controller, see Fig. 1 right). To stop placement, the same button is pressed again (counts/landmarks each require a single press). The relation between successive points is implied, that is, no segments are drawn by default, unless explicitly activated by the user to serve as visual aids. The point tool thus by design trades contouring accuracy for shorter delineation times. By minimising button pressing and keeping the user engaged continuously placing points, a sense of flow may be promoted [2, 3]. To adapt performance to the data at hand, the duration of the time interval is configurable and the pointing source can be switched to follow the user’s gaze. Gaze interaction has been designed to stop as soon as the user looks away.

#### Line tool

A second tool, called the line tool, addresses the drawing of segments in conjunction with the placement of points. In contrast to the point tool, the line tool trades longer delineation times for higher contouring accuracy. With it the user draws segments by also placing points successively, but segments are interpolated using natural cubic splines [23], where the placed points serve as control points. To close a sequence of segments, the user must “click” in the vicinity of a *start point*. Unlike regular points, start points are interactive and react to hovering. Additionally, each segment is annotated with a user-oriented, equidistantly placed label from the segment’s ends that indicates its anatomical length.

#### Underlying data model

To formalise how points and segments are stored for quantification, we implemented a graph-based data model. That is, after image data has been interactively annotated by placing points and segments, these are added to the data model. In the data model, points and segments comprise nodes and edges, respectively, such that any node has a maximum degree of two. Depending on the delineation tool, a node *n* can be represented as an isolated point (*deg*(*n*) = 0), as a start/end point in a spline (*deg*(*n*) = 1), or as an intermediate point connected to two other points (*deg*(*n*) = 2). For simplicity, only the nodes are provided directly by the user, the edges link nodes either as drawn segments derived from the nodes or as implied edges not shown to the user. In the model, every node belongs to a group, which is referred to as *region*, such that delineations are grouped to correspond to different anatomical areas, such as brain regions. All other elements and annotations are derived from the points and segments.

#### Interactive editing

As part of the VR-editor functionality, delineations can be interactively edited using a number of dedicated interactions and interfaces. The design of those has been based on established metaphors, such as those on the desktop, in order to be easy to use, including for users who may not be familiar with VR. One such interaction based on a familiar metaphor is 3D selection, where selection is realised as a natural extension to 3D from the controller’s input. As the user moves the controller, the operation incrementally updates a 3D envelope, where any points falling inside this are marked as selected 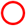. Further operations can then be performed on the selected points, e. g., deletion and region re-assignment, which are grouped in a dedicated VR context menu. Using the menu, for example, a region comprising a set of points can be created, deleted, selected, assigned elements to, and shown or hidden.

For quick editing, undo functionality is also provided. An undo is triggered with a swipe in the air with the off-hand controller, horizontally to the left of the user 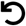. To avoid accidental triggering, the swiping interaction is modified by requiring the user to hold the controller’s grip button, which is otherwise used for direct object manipulation.

#### Interaction aids

To support the interactive delineation in VR, we designed several additional interaction aids [4, 5]. While moving around in VR, text labels may become illegible due to their fixed orientation, while continuous reorientation may be disturbing. Our solution, therefore, was to couple reorientation with a concurrent event that is of greater saliency. The design reorients annotations by exploiting certain occurrences of inattentional blindness [24], such as during point placing by the user. Beside annotations, tissue may also be imaged in a way that disagrees with the user’s mental model. For this, the imaged tissue can be re-aligned by interactively rotating its voxels, with rotation constrained to multiples of 5^*◦*^ to support reproducibility. Additionally, if the user decides to increase the object scale of the tissue in VR, they can so implicitly configure delineation either for higher precision (larger scale) or for shorter delineation times (smaller scale). Thus, larger scales could not only be used to strengthen immersion [25], but also as a parameter for adjusting precision.

Direct manipulations are also constrained to reduce associated costs, such as for picking up objects. Constraining interaction is also important for supporting more reproducible delineations across multiple datasets. For example, orthoplanes can be interactively shifted to map onto single images from the image stack or advanced at regular intervals, multiples of 5 images. In this way, a user can shift orthoplanes uninterruptedly, but always stop at a single image slice. Lastly as a necessary condition for reproducibility, delineation data can be saved and shared natively or exported as coordinates to other software tools and frameworks [9, 10]. By using hidden signatures that are interoperable with other software tools, coordinate values may be fully re-imported as points and line segments. Fig. 2 illustrates this workflow.

**Fig 2.**
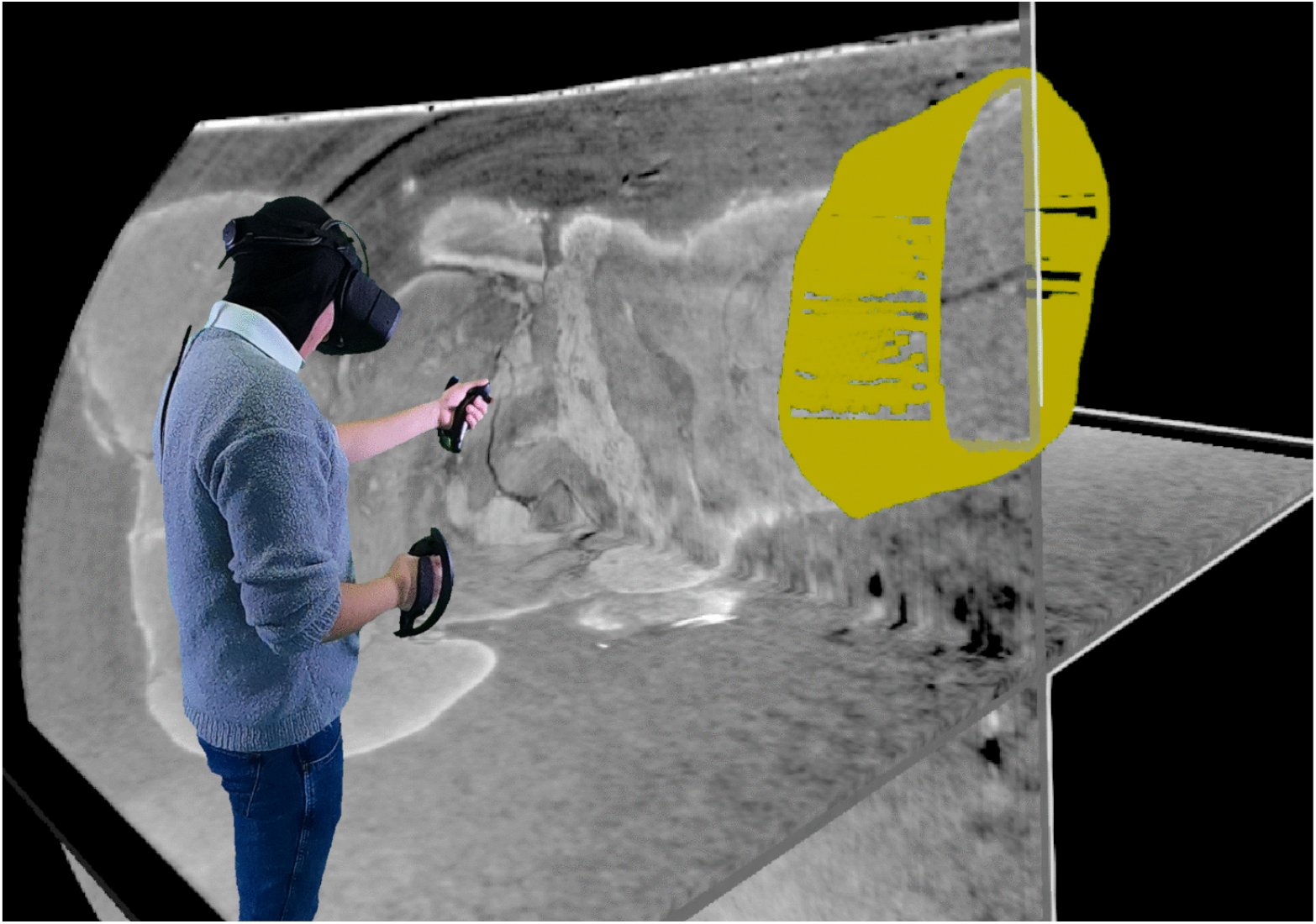
A user explores delineations inside VR which have been created on the desktop. The delineations, shown in yellow, cover part of the brain stem of the imaged animal. The image illustrates how *Brainacle* could be used with other software tools, in this case Fiji [10], for the purposes of VR exploration.

### Quantification of structural features

Cichlid gross brain measurements commonly comprise the length, width, depth, and weight [26] or the volume and centre (*x, y, z*)-positions [27]. The measurements that may be performed often depend on the imaging modality that has been used to image the data. For example, histological sections, have high spatial resolution (e. g., allowing to determine cell densities), but are less suitable for volume measurements. Further along the pipeline, measurements are also affected by how image data has been 3D reconstructed. For example, in the case of isosurface meshes measurements are derived from higher-order polygons (such as triangles), which may affect precision downstream. For this, we considered measurement methods that do not require a prior surface reconstruction in the image space and instead rely on user input in the object space. We tested the methods’ calibration, i. e., that any performed measurements are anatomically correct, by performing validation inversely: predefined positions were read and measurement values were then filled. Using a single dimension for simplicity, we set positions for which the ground truth is known and compared the result after calculation.

#### Slice area

We apply Lebesgue integration to measure the area of a delineated slice in order to derive the volume of a region from the sequence of delineated slices. Unlike voxel-counting, such as in the heart [28], in the brain [29], or their implementations in available software [9, 10], integration generalises this to derive area measurements from a sparser user input in the object space, while still allowing for a configurable level of accuracy and computational cost. As a result, we propose the `SliceArea` algorithm that calculates the area of a single delineated slice as illustrated in Fig. 3.

**Fig 3.**
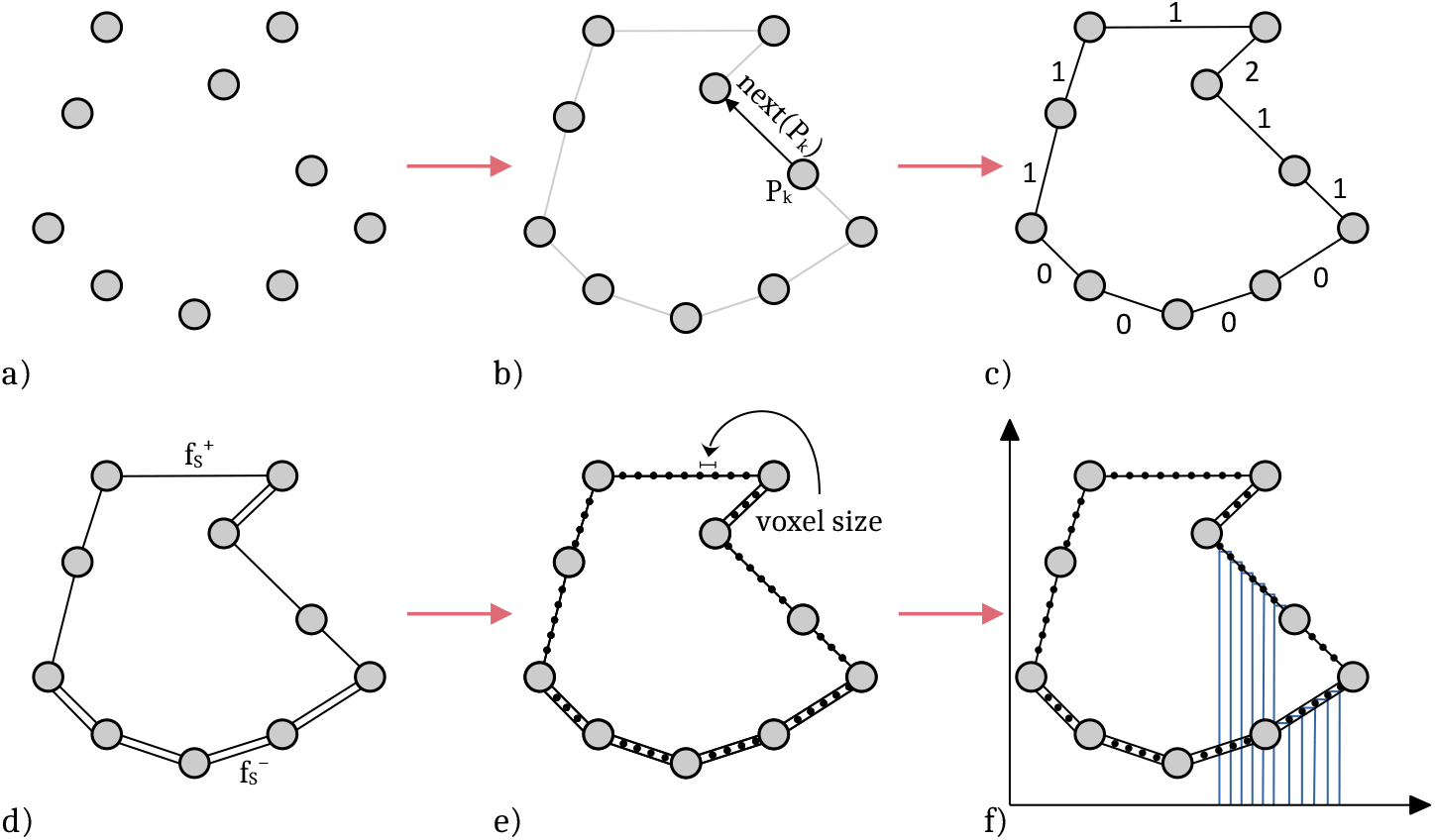
Slice area. a) A set of coplanar points, b) Connections (edges) are defined a priori, c) Outer (even) and inner (odd) edges are determined using the number of intersections of the rejections from *X* to points on the edge, with all other edges. d) Outer edges contribute signed negatively, while inner edges signed positively. e) Intra-segment points are interpolated using the voxel size to construct 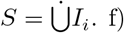) Each function is Lebesgue-integrated over *S* with respect to *µ*, where *µ*(*I*) is measured as the *e*_*x*_-norm of *I*.

##### Algorithm 1

Slice Area

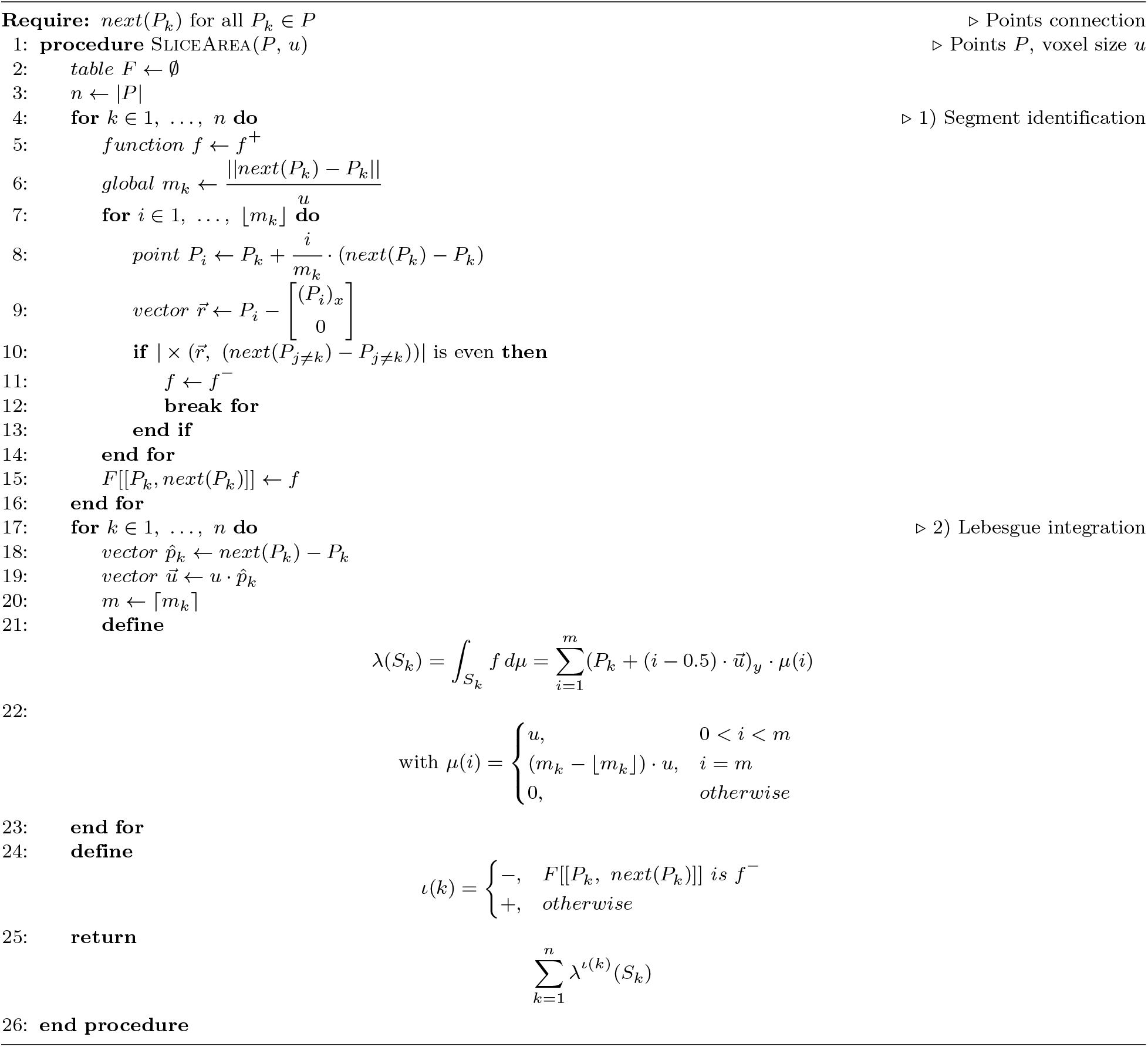

The algorithm, given in Algorithm 1, consists of two phases: 1) Segment identification and 2) Lebesgue integration. If the segment positions are specified (i. e., drawn with the line tool), these are treated as points by the algorithm. Otherwise, the only points used as input by the algorithm are the points placed by the user. In each case, the interpolation is linear. Segment identification is described in Fig. 3 c)–d). Lebesgue integration is performed over the collection *L*(*S, S, µ*), where *L* describes a slice, consisting of measure spaces (*S, S*, *µ*)_*k*_ with *k* = 1, …, *P* for *P* the set of points. The set *S* represents a segment, is its *σ*-algebra, and *µ* : *S* → ℝ the measure on *S*. For an intra-segment *I* ⊂ *S* with *µ*(*I*), we select *u*, the voxel size, such that *µ*(*I*) ≤ *u*. In general, the collection of Lebesgue-measurable spaces *L* may be disjoint; connectedness is sufficient, but not necessary, and correctness depends on how point connections are defined.

Algorithm 1 expects points to be coplanar and it requires a function *next*(*P*_*k*_) that supplies the next point from a point *P*_*k*_ on the *k*^th^ slice. At line 1, the procedure SliceArea has two parameters: a set of points *P* ⊂ ℝ^3^ and a voxel size *u*. Segment identification (lines 4–16) proceeds by determining the number of measures for each segment (line 6). Then, the rejection 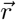 from the *X* axis to each point *P*_*i*_ (line 8) is determined (line 9). If an even number of intersections between 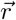 and any other segment is encountered (line 10), the positive part *f* ^+^ of the simple function *f* (line 5) is replaced with the negative part *f*^−^ and execution for the current segment stops (lines 11–14). The negative part *f*^−^ is negatively signed to correspond to an outer segment, the values of which are subtracted (see Fig. 3 f)). At line 15, *f* is stored in table *F* for further processing in step 2) Lebesgue integration. At lines 18–20, several working variables are introduced for the calculation of the intra-segment values of *f*. The Lebesgue integral of the simple measurable function *f* over a segment *S* with respect to *µ* is defined as the sum of the products of the intra-segment values and their measures *µ*, which are defined on line 22. The measure of a segment *λ* is signed, with its sign determined using table *F* (line 24). The sum of all signed measures of all segments returns the slice area (line 25), completing the procedure.

#### Slice volume

Now, it suffices to extrude the measure using the voxel size to obtain the volume of a single slice: *µ*_2_ : *S* → ℝ, *µ*_2_ (*I*) → *µ*(*I*) · *u*, and

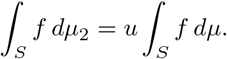

Then, the sum over all such slices results in the region volume:

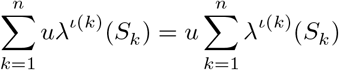

In the case of intermediate non-delineated slices, the computation requires slice interpolation, that is, at the length of the voxel size *u*. An axis-aligned rotation of the data may be performed if necessary.

#### Other shape measures

Other shape measures are provided to aid the analysis of delineated regions. These include the region’s centroid, its convex hull volume [30], the region’s major, minor and intermediate axes, as well as the volume of the minimum-volume enclosing ellipsoid [31]. Anatomical ratios may be inferred by interactively drawing with the line tool, while the region’s convexity and compactness can be derived from the ratio of the actual to the convex hull volume and to the ellipsoid volume, respectively [32].

## Results

Measuring brain size has long been a core aspect of many approaches in neuroethology, in particular in studies examining the social brain hypothesis—a controversial concept which posits that larger brains have been selected for by increased cognitive demands of living in groups [33]. However, recent work suggests overall brain volume is a poor indicator of underlying cognitive processes [34], and that regional specialisation may be a better predictor of cognitive performance than size per se [35]. The relative size and connectivity among brain regions may be more meaningful measures than gross size [36], demonstrating that researchers require more sophisticated tools to measure regional volumes in the brain. The precise delineation of brain regions and substructures therefore constitutes a foundational step in establishing new model systems for studying social evolution.

To this end, *Brainacle* was applied to a case study in the domain of neuroethology, thereby validating its functionality through longitudinal use. A demonstration of the obtained results is shown in Fig. 1. Case-study demonstrations are often employed when new desktop bioinformatics systems and pipelines are first proposed, in contrast to new VR systems which are often required to be evaluated in a laboratory user study. User studies of new systems, however, may suffer from novelty biases [37] and high preference and usability in the lab do not guarantee lasting usefulness for the community [38, Tab. 1]. Such system-specific evaluations are also less likely to be independently replicated by other researchers, whereas applying a tool to a use case merely validates the tool’s capabilities for that use case. This serves to communicate the tool’s functionality to other researchers and it does not involve making statistical claims about the respective user population (e. g., *biologists find this tool to be user-friendly*), that is, claims relating to a specific version of the software tool which may not be up to date any longer.

### Tissue reconstruction

Tissue reconstruction followed the processing pipeline shown in Fig. 4. All species comprised lab-reared cichlids from the tribe *Lamprologinii*, endemic to Lake Tanganyika, Africa, and naturally occurring in Zambia: *Neolamprologus multifasciatus, Neolamprologus brevis, Lamprologus ocellatus, Lamprologus callipterus, Telmatochromis temporalis*, and *Lamprologus ornatipinnis*. According to standard ethical procedures, fish were euthanized in a water bath of MS-222 dissolved in holding water (0.50 mg/mL) and subsequently rapidly decapitated to ensure death. Brains were carefully removed and stored over night in 4% formalin solution in PBS. Afterwards, the brains were placed in tissue embedding cassettes (Simport Scientific Macrosette™) and dehydrated in an ascending ethanol series before being embedded in paraffin (Leica Histowax, melting point 57-58^*◦*^C). The brains were imaged at the SYRMEP beamline (SYnchrotron Radiation for MEdical Physics) of the Italian synchrotron “Elettra” in Trieste, Italy. The beamline was to perform propagation-based imaging in white beam mode [39]. We then used *Syrmep Tomo Project* (STP) [40] v1.5.3 to obtain 32-bit floating point image sequences, where reconstruction comprised of pre-processing with conventional flat-fielding and de-stripping [41], Paganin *et al*.’s phase retrieval [42], and filtered back projection. For the phase retrieval, the pixel size was 2.016 *µm*, the distance was 200.0 *mm*, and the energy was 25.0 *keV*; the delta-to-beta ratio was *δ/β* = 300. We provide the full protocol [8] of the pipeline that is shown in Fig. 4.

**Fig 4.**
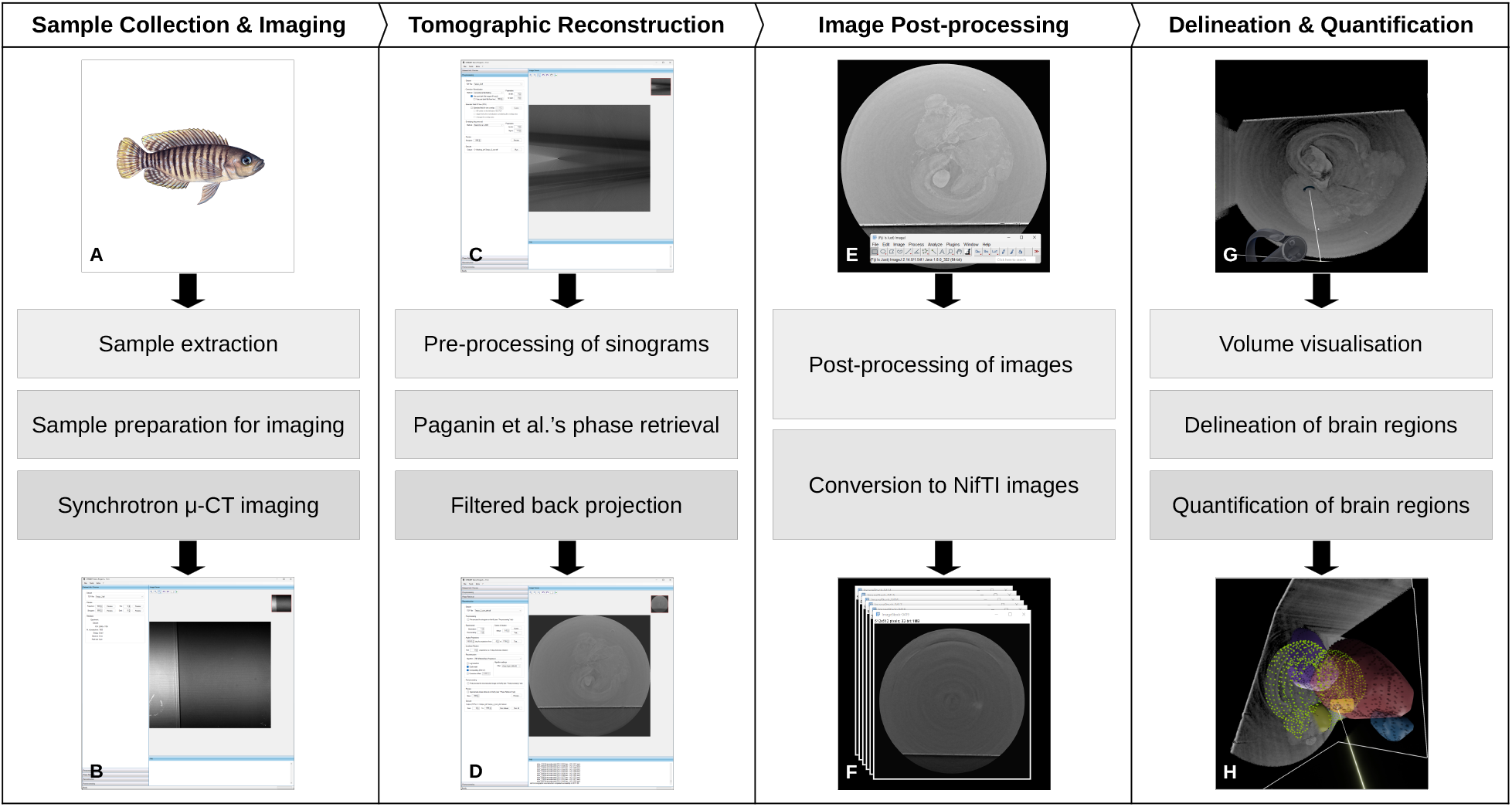
Processing pipeline. Left to right, the four stages are 1) sample collection & imaging, 2) tomographic reconstruction, 3) image post-processing, 4) delineation & quantification. Each stage has an input and output: A) cichlid of the species *N. multifasciatus*; B) preview of the central projection in STP [40]; C) preview of the central sinogram in STP; D) preview of the central slice after reconstruction in STP; E) preview in Fiji [10], with contrast enhancement of 0.1%; F) image stack after down-scaling and conversion to NIfTI in Fiji; G) a view in *Brainacle*; H) delineated gross brain regions in *Brainacle*.

### Delineation

To set the voxel size in *Brainacle*, a file of the same name but a different ending was created in the directory of the image dataset. This contained the voxel size with its SI unit of length, i. e., 8.064 *µm* accounting for the dataset’s change in scale. Once loaded into *Brainacle*, each dataset’s voxels were rotated in VR and their rotation was noted down. Voxel rotation was used to align the user’s current view in VR with a textbook view of the brain, i. e., from the posterior to the anterior. During delineation, brain atlases of related species, such as tilapia [27] and zebrafish [43], were consulted and brain regions with tearing or blurring of the tissue were excluded. Slices were delineated using the point tool, for which the analyst advanced the delineation plane at regular intervals from the anterior to the posterior of the brain (every 5^th^ slice). The following gross brain regions were delineated for a given image dataset: olfactory bulbs, telencephalon, optic tectum, hypothalamus (and hypophysis), and cerebellum. The completed delineations, in total amounting to about 100 delineated brain regions, were then saved in order for quantification, the next step in the pipeline, to proceed separately.

### Quantification

Several measures were computed using *Brainacle* for each of the delineated brain regions, these are aggregated in Tab. 1. A minimum-volume bounding ellipsoid was fit to obtain the length of the principal axes of each delineated brain region. Brain region volumes were measured using two different approximations: a convex hull and a minimum-volume bounding ellipsoid. Using the annotations in VR, region IDs were then manually mapped onto region names for further statistical analysis using R.

To compare the convex-hull volume to our implementation [31] of the classical ellipsoid method known from whole-brain photography, we correlated the two measures by brain region. Using box plots, one outlier, for olfactory bulbs, was detected, which might have been due to the region’s relatively smaller size. The convex-hull and bounding-ellipsoid measures were highly correlated across all brain regions with *r* coefficients upwards of 0.97, indicating similar performance (see the figure in our research dataset [8]). Notably, the bounding-ellipsoid volume always overestimated the convex-hull volume. These observations point to two conclusions in the context of interspecies comparison. First, more precise but also interpretable measurements, e. g., Alg. 1, have the potential to reduce some of the uncertainty associated with approximation measures, which continue to find wide application due to their relative simplicity. Second, such results also corroborate interactive VR as an alternative method for delineating and quantifying anatomical structures. To show that a non-trivial number of brain regions (ca. 100) can be interactively delineated using VR, we used the absolute brain sizes to report the aggregate measurements in Tab. 1. For further biological interpretation and analysis, relative brain sizes have to be obtained to account for the individual variations in body size.

## Discussion

In the presence of already widely used and affordable image-analysis software for the desktop [9–12], immersive and virtual environments, such as those in VR, should target well-defined problems and applications [1], with justification given by the affordances of those environments [2, 4, 5]. As one such application, tissue delineation represents a visuospatial task that requires understanding anatomical structure at the level of the individual animal. VR, with its embodied navigation and 3D visualisation, can support this task by allowing users to quickly gain an overview of the relevant anatomy and interactively add delineations inside the same 3D environment. A native VR editor can also be applied to new or diverse data with relative ease. Users may achieve this by visualising the data without specialised preprocessing before utilising VR’s interactivity and gamification to delineate this. For example, the lack of tissue staining was not an issue for the delineation in VR, in contrast to our initial approach which relied on standard image operations from desktop software in order to create a binary mask. Furthermore, by allowing the user to control the scale of the visualisation, users can adapt the level of precision as necessary during delineation.

With the case study of African cichlids, we have demonstrated that VR can be used to delineate and quantify a substantial number of different anatomical structures. In the same time as this class of VR software tools holds promise to be applicable more generally, certain methodological limitations also became evident in the process of designing and iterating *Brainacle*. A distinction between efficient and exact quantification emerged based on the specific delineation. On the one hand, good approximations could be readily obtained compared to classical methods using whole-brain photographs and histological-sectioning pipelines. On the other hand, exact quantification still requires additional considerations. Indeterminate values for *next*(*P*_*k*_), prevented us from applying Alg. 1 to the set of delineations without additional region assignments. Similarly, in the case of multiple or disconnected substructures, separate regions should be assigned to each substructure or both delineation tools should be used to indicate where each substructure begins and ends. There are also well-known limitations to using VR. For instance, for the purposes of usability we had to minimise users’ text entry inside VR by requiring text values to be configured before entering VR. And while we paid considerable attention to enable the support of replicable results using *Brainacle*, e. g., by introducing constrained interaction in VR and serialising additional data from the environment, delineations may still be user-dependent. It remains an open question for future research to determine how strongly quantifications may be influenced by variations in the input data. In the meantime, however, new software tools and hardware setups could provide researchers with greater optionality and broaden the scope of existing applications.

In conclusion, we have presented *Brainacle*, a VR editor for the interactive delineation and quantification of anatomical structures that can be applicable to different biological pipelines. *Brainacle* has been developed in an effort to encapsulate and make our workflow available to the wider research community. User documentation about using the software is available on the website https://brainacle.org, while research data for the case study has been deposited for long-term preservation on the data repository DaRUS [8].

## Data availability and ethics

*Brainacle* runs on various HMDs and controllers connected to a Windows operating system, including prior and current generations of Oculus and Meta Quest (via Quest Link), Valve Index, Pico Neo (via Streaming), HTC Vive, Oculus Rift, Varjo, and Pimax. It is available at https://brainacle.org and on DaRUS [8] for use free of charge. Please refer to the user license agreement at https://doi.org/10.18419/DARUS-4779. Except for an example dataset [8], the image data and absolute measurements are part of ongoing research, but are available from Alex Jordan on reasonable request.

For development of *Brainacle* we used Unity, in addition to the following libraries. *Vive Input Utility* and Valve’s *SteamVR* were used and extended to provide VR input. Volume data is rendered using LISCINTEC’s Volume Viewer, a commercial software package. To generate convex hull isosurfaces, we integrated the *MIConvexHull* software package. For spline interpolation, we used *TinySpline* [23]. *Unity Color Picker* and *Unity Standalone File Browser* were also used as common Unity dependencies.

Brains were collected according to Germany’s Tierschutzgesetz, BGBl. I 2013 S. 2182 and the EU’s Directive 2010/63/EU. The euthanasia of fish for extraction of brains was registered under file number T-17/11TFA at the responsible state authority. No procedures were conducted on the animals, no behavioural or other data was collected. The study followed the ARRIVE 2.0 guidelines for animal research. The persons demonstrating the software tool in Fig. 1 and Fig. 2, respectively, are two of the authors.

## Acknowledgments

The authors thank Bärbel Heidrich for her input during imaging at Elettra, Jinman Kim for his feedback about the software, Francesco Brun for his advice regarding STP, and Sylvia Garza Reyes regarding prior image processing. This work was supported by Deutsche Forschungsgemeinschaft (DFG, German Research Foundation) under Germany’s Excellence Strategy—EXC 2117—422037984, as part of SFB/Transregio 161—DFG Project ID—251654672, and by the Max Planck Institute of Animal Behavior. Beamline time and technical support for imaging were supported from Elettra under proposal 20170091 for the SYRMEP beamline.

